# GeneSqueeze: A Novel Lossless, Reference-Free Compression Algorithm for FASTQ/A Files

**DOI:** 10.1101/2024.03.21.586111

**Authors:** Foad Nazari, Sneh Patel, Melissa LaRocca, Ryan Czarny, Giana Schena, Emma K. Murray

## Abstract

As sequencing becomes more accessible, there is an acute need for novel compression methods to efficiently store this data. Omics technologies can enhance biomedical research and individualize patient care, but they demand immense storage capabilities, especially when applied to longitudinal studies. Addressing the storage challenges posed by these technologies is crucial for omics technologies to achieve their full potential. We present a novel lossless, reference-free compression algorithm, GeneSqueeze, that leverages the patterns inherent in the underlying components of FASTQ files (*i*.*e*., nucleotide sequences, quality scores and read identifiers). GeneSqueeze provides several benefits, including an auto-tuning compression protocol based on each sample’s distribution, lossless preservation of IUPAC nucleotides and read identifiers, and unrestricted FASTQ/A file attributes (*i*.*e*., read length, read depth, or read identifier format). We compared GeneSqueeze to the general-purpose compressor, gzip, and to the domain-specific compressor, SPRING. GeneSqueeze achieved up to three times higher compression ratios as compared to gzip, regardless of read length, read depth, or file size. GeneSqueeze achieved 100% lossless compression, with the original and decompressed files perfectly matching for all tested samples, preserving read identifiers, quality scores, and IUPAC nucleotides, in contrast to SPRING. Overall, GeneSqueeze represents a competitive and specialized compression method optimized for FASTQ/A files containing nucleotide sequences that has the potential to significantly reduce the storage and transmission costs associated with large omics datasets without sacrificing data integrity.

## Introduction

The field of genomics, due to advancements in high-throughput sequencing technologies, has experienced a revolution in the past two decades, with almost a million-fold reduction (from 1 billion to hundreds of dollars) in the cost of sequencing.^1^ The increase in accessibility has led to an increase in the quantity of omics data, with estimations predicting that 2–40 exabytes of data will be generated within the next decade.^2^ Unfortunately, storage technology is advancing at a much slower pace, leading to critical technical and economic bottlenecks for omics-based discovery and their applications in biomedical or clinical research. Omics data is essential in personalized medicine which relies on accurate omics data to extract biomarkers or signatures that can individualize a patient’s preventative measures, diagnoses, treatments, and monitoring.^3^ To be practically used in patient care, omics data needs to be stored, retrieved, and transmitted in a manner that is cost-effective, time-efficient, and lossless for analysis and dissemination. Therefore, specialized compressors for omics data are needed to continue to support advancements in medicine.

Nucleotide-based omics data (*i*.*e*., genomics, transcriptomics, epigenomics) are commonly stored in FASTQ or FASTA text-based formats. FASTA is a simple format that stores only the read identifier lines followed by nucleotide or amino acid sequences. FASTQ files are more complex, storing information on nucleotide sequences as well as the quality scores for each base. The addition of quality scores makes FASTQ the preferred format for sequencing platforms (*e*.*g*., Illumina) and downstream applications as they reflect the level of confidence in the read of each base in a sequence.^3^ Quality scores are critical for analyses such as variant detection, where the accuracy and reliability of the base calls are foundational.^4,5^ Both FASTA and FASTQ files can be large, particularly for high-coverage sequencing data, which makes them difficult to store and process efficiently. Therefore, FASTQ/A files are routinely compressed for more practical storage, management, and transmission.

Presently, biologists are employing general-purpose compression tools. These tools are not tailored specifically for biological data and thus do not fully leverage the inherent repetitive characteristics of FASTQ files, leading to lower compression efficiency. Gzip^6^, a general-purpose algorithm that combines Huffman encoding^7^ and LZ77^8^ to create a dictionary tailored to the frequency of word repetitions in the data, performs relatively poorly on genomic data compression. Despite the inefficiencies, gzip remains as the de facto standard in the biology domain due to its stability and popularity. Gzipped files are accepted as input by various sequencing analysis tools and are used by public repositories of genomic data.^9^ Alternatively, domain-specific algorithms that leverage the redundancy in FASTQ files, and explicitly utilize the underlying structures present in the nucleotide reads, quality scores and read identifiers of genomic data, have shown promise in genomic data compression, achieving high compression ratios while maintaining high accuracy for downstream analysis.

These domain-specific FASTQ compressors utilize underlying compression methodologies such as classical, read-mapping-based, read-assembly-based, or read-reordering-based.^10^ Among these, classical compressors are notably less efficient. Unlike the other three categories, which necessitate auxiliary information such as a reference sequence or preprocessing steps like sequencing reordering prior to compression, classical compression methods operate solely on reads in their original form and lack the ability to leverage read redundancies.^10^ Read-mapping-based methods map nucleotide sequences to a provided reference genome, and encode the alignment position and the probable mismatches.^10,11^ Read-assembly-based compressors encode nucleotide sequences by using built-in long contiguous sequences (contigs), as opposed to a reference genome.^10,12^ Alternatively, read-reordering-based compression methods are reference-free and involve arranging overlapping nucleotide sequences near each other, to encode reads more efficently.^10,13^

Various domain-specific genomic data compressors such as FaStore^10^, SPRING^14^, FQSqueezer^15^, PetaGene^16^, Genozip^17^, ColoRd^18^, repaq^19^, MZPAQ^20^ have shown significant promise in addressing data size challenges within the domain. In our study we compare our algorithm to the domain specific compressor, SPRING, a compressor which combines read-assembly-based and read-reordering-based methods. SPRING is based on the HARC algorithm and supports pair-preserving compression, lossless compression (with the exception of certain cases such as FASTQ/A files containing rare IUPAC nucleotides or specific read identifier formats), and lossy compression of quality values, among other features.^14,21^ Unfortunately, each of the existing domain-specific compression methods, including SPRING, still have disadvantages, including poor compression ratio, lossy compression, nucleotide sequence - quality score-encoding dependency, limitations on long reads, poor performance for nucleotide sequences that include N bases, lossiness for non-ACTGN IUPAC bases, and/or rigid protocols.

With the varying weaknesses of existing compression methods, there is a need for efficient, domain-specific methods for next-generation sequencing (NGS) data compression. We present a novel reference-free compressor, GeneSqueeze, which uses read-reordering-based compression methodology. Notably, the reordering mechanism within GeneSqueeze is temporally confined to specific algorithmic blocks, facilitating targeted operations on the reordered sequences. Upon completion of these operations, the residual sequences are reconstituted to their original order, thereby preserving losslessness without necessitating the retention of the original index. GeneSqueeze presents a dynamic protocol for maximally compressing diverse FASTQ/A files containing any IUPAC nucleotides while maintaining complete data integrity of all components, verified by MD5^22^ matching.

## Methods

### FASTQ/A format

The representation of data obtained from a sequencing experiment follows the FASTQ format introduced by Cock *et al*.^23^ Each read and its corresponding metadata are encapsulated in a block of four lines. The read identifier is present on the first line and provides information on the sequencing run. The second line contains the nucleotide sequences (A, C, T, G, N) that are called by the sequencer. In most FASTQ/A files, among non-ACGT (irregular) bases, only N is common, and the other irregular nucleotides identified in IUPAC standards presented in **Table 1**. are rarely observed. The fourth line contains the quality score for each nucleotide in the read, which indicate the probability that each base in the read is incorrect. The quality scores are encoded in ASCII using Phred scores.^23^ The third line contains the symbol ‘+’ to separate the nucleotide sequences and the quality values, and may also include metadata or comments. An example of the first eight lines of a FASTQ file is shown in **Fig. 1**.

**Table 1.** IUPAC (International Union of Pure and Applied Chemistry) single letter codes for DNA/RNA.

| Symbols | Name | Remarks |
| --- | --- | --- |
| A | Adenine | Purine |
| G | Guanine | Purine |
| C | Cytosine | Pyrimidine |
| T | Thymine | Pyrimidine |
| U | Uracil | Pyrimidine |
| R | Purine | A or G |
| Y | Pyrimidine | C or T/U |
| M |  | A or C |
| K |  | G or T |
| S | Strong | C or G |
| W | Weak | A or T |
| H | Not G | A or C or T |
| B | Not A | C or G or T |
| V | Not U/T | A or C or G |
| D | Not C | A or G or T |
| N | Ambiguous | A or C or G or T |

**Figure 1.**
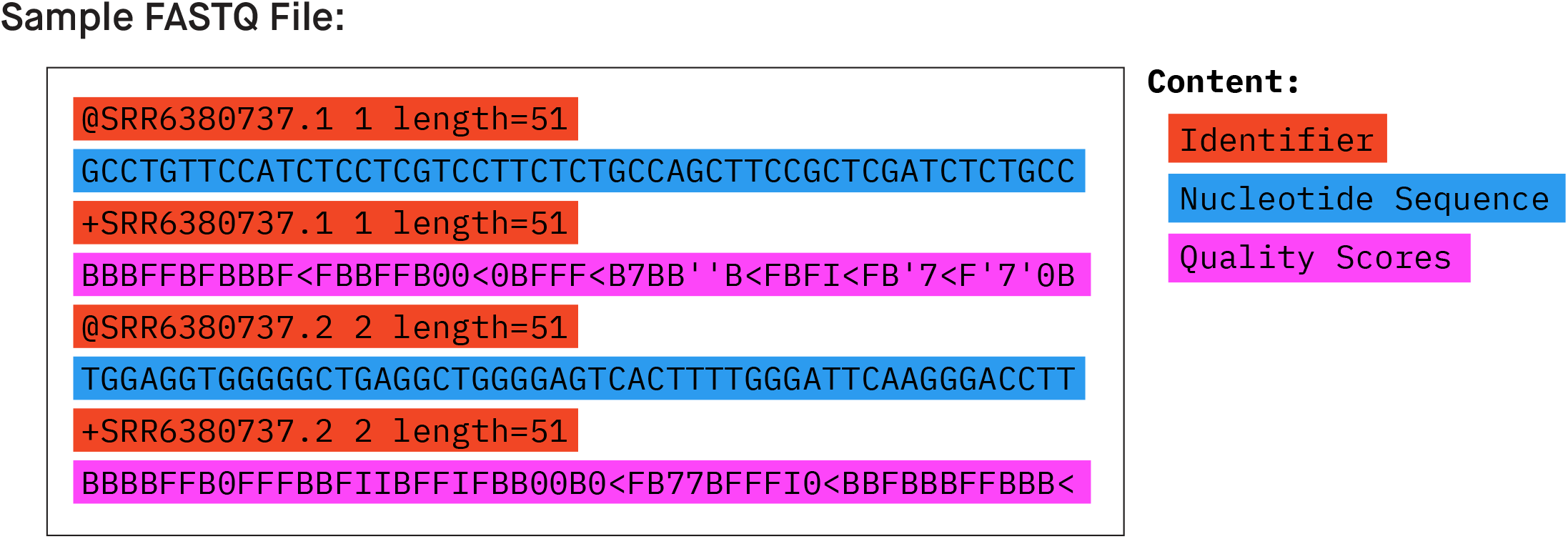
An example FASTQ file

### GeneSqueeze Algorithm

In this section, the proposed algorithm for compression of FASTQ/A data is described. The algorithm separates the nucleotide, quality score, and read identifier sequences and then passes each sequence type through a specific branch of the GeneSqueeze algorithm, as presented in the flow diagram in **Fig. 2**. The way this algorithm both branches and blocks function is explained in this section.

**Figure 2.**
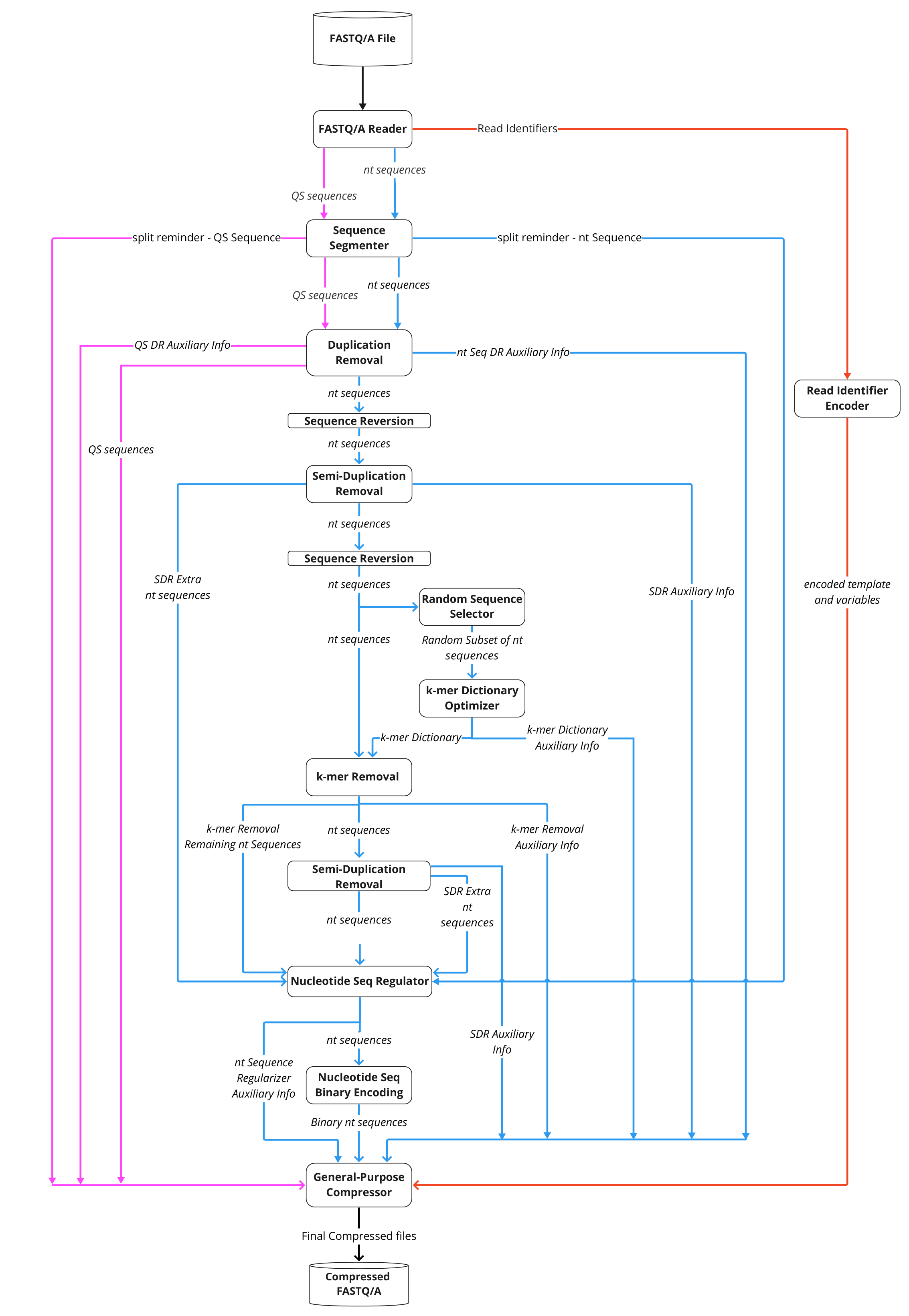
GeneSqueeze Compression Flow-Diagram

#### FASTQ/A Reader

The *FASTQ/A Reader* block loads and reads the FASTQ/A files to memory before splitting the file into nucleotide sequences, read identifiers, quality score sequences (for FASTQ only).

#### Sequence Segmenter

The *Sequence Segmenter* splits long nucleotide sequences and quality score sequences into smaller segments to improve overall compression efficiency. Our study found that there exists an ideal sequence length (*ideal_len*) to maximize GeneSequeeze compression efficiency. In this version of GeneSqueeze, a fixed value for ideal length was assumed for all samples/datasets. The number and length of created segments from each nucleotide and quality score sequence is determined using equations 1 and 2, respectively. The floor function, denoted as ⌊x⌋ and employed in equations 1 and 2, yields the greatest integer less than or equal to x.

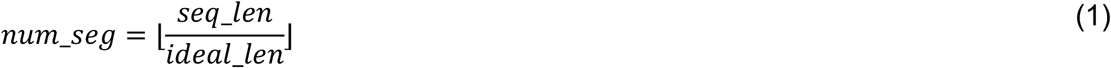

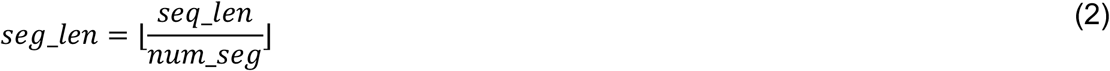

Where *seq_len, seg_len* and *num_seg* are the length of input sequences, length of created segments, and number of segments, respectively, and the *ideal_len* is a hyperparameter that is used to find the *seg_len*.

The next block of the algorithm, *Duplication Removal*, necessitates equal segment lengths are present for processing of nucleotide and quality score sequences, and thus, if (*seq_len* > *num_seg* × *seg_len*), the remainder (*extra_len* bases) of reads are separated and concatenated together to make a long sequence to be processed separately.

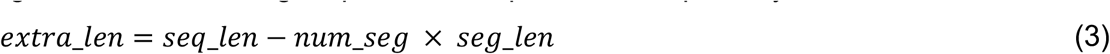

Following is a numerical example:

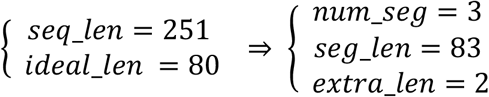

If the input to the *Sequence Segmenter* is a nucleotide sequence, the remainder sequence is directed to the *Nucleotide Sequence Regulator* block; if the input is quality score sequence, the remainder goes to the *General-Purpose Compressor* block.

#### Nucleotide Compression Branch

The segmented nucleotide sequences are passed to the nucleotide sequence compression branch of the algorithm, which consists of the following blocks: *Duplication Removal* (DR), *Sequence Reversion, Random Sequence Selector, k-mer Dictionary Optimizer, k-mer Removal, Semi-Duplication Removal (SDR), Nucleotide Sequence Regulator, and Nucleotide Sequence Binary Encoding*.

#### Duplication Removal (Nucleotide Branch)

The main objective of the *Duplication Removal* block is to find, group, encode and remove all duplicate reads in an efficient manner (example in **Fig 3**). The block first creates a temporary index, called original index, for each sequence to preserve a reference to the original sequence order, during the sorting of the sequences alphabetically. Identical sequences are then grouped. The first instance of the nucleotide sequence is labeled as the parent sequence and all subsequent instances are designated as child sequences. An identifier label (*dr_identifier*) stores information on parent and child sequences. If the sequence is denoted as a parent sequence, the algorithm then identifies if the parent sequence has any child sequences, before labeling them accordingly. If the sequence is a child sequence the identifier stores the index of its parent sequence so the child’s sequence can be retrieved during decompression. The child sequences are then removed and the parent sequences are re-sorted using their original index, and the temporary original index value is removed.

**Figure 3.**
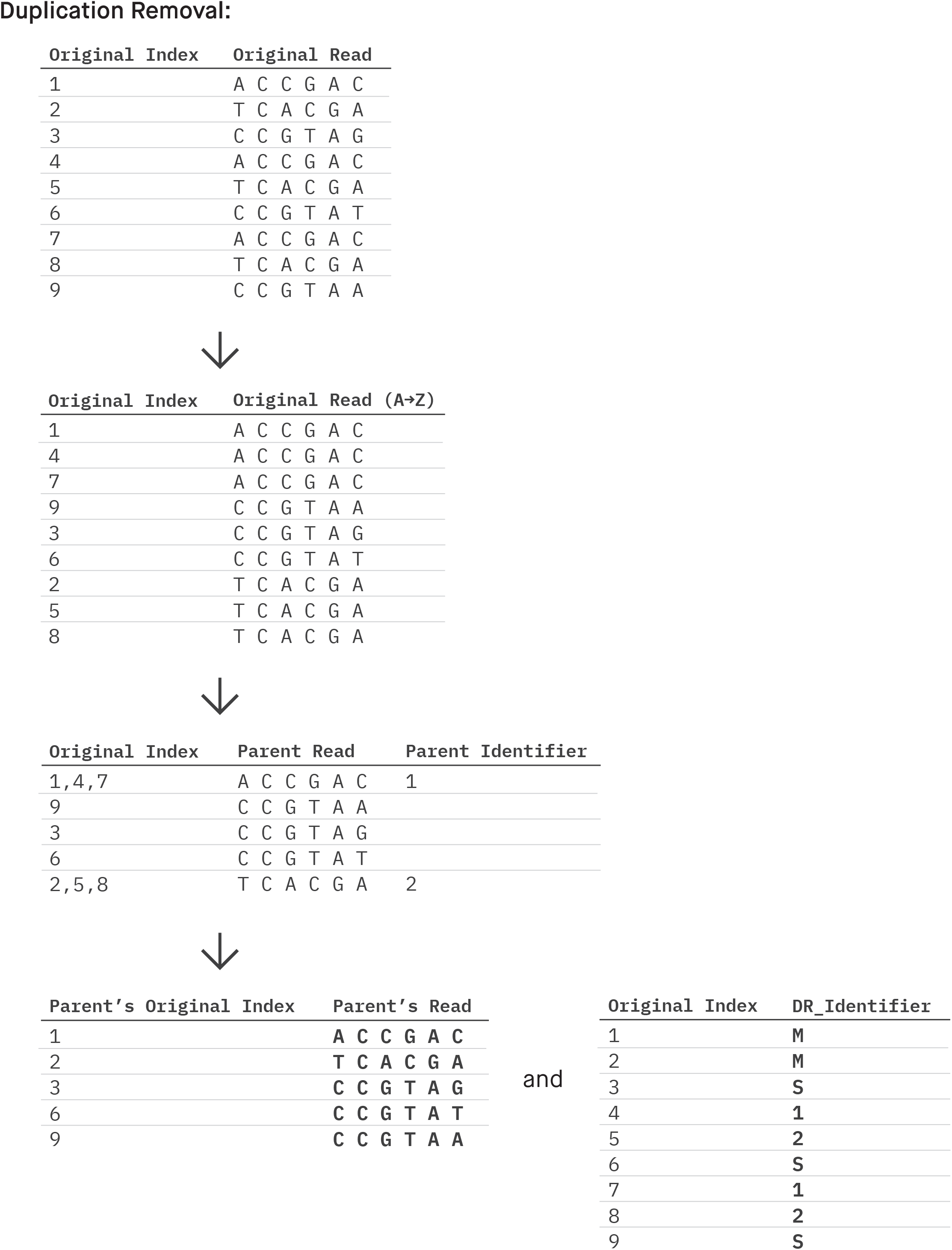
GeneSqueeze Duplication removal process

The pseudocode presented below shows the duplication removal process: FUNCTION duplication_removal(sequences)

1. Convert sequences to an array
2. Sort the sequences and keep track of their original indices
3. Group duplicate sequences and maintain original index values
4. Identify ‘parent’ sequence as first sequence of each group
5. Identify ‘parent’ index as index of each group
6. Prepare two data structures to store information about sequences and groups One for the parent sequences One for identifier label (dr_identifier)
7. Parent sequences are placed into a DataFrame and sorted based on original index
8. Each original index is assigned an identifier in dr_identifier—-IF Index is associated with a parent sequence with at least one child THEN equals ‘M’ ELSE IF Index is associated with a parent sequence with no child THEN equals ‘S’ ELSE IF Index is associated with a child sequence THEN equals its parent index
9. Return the sorted sequences and the dr_identifier

The parent sequences are passed to the *Sequence Reversion* block and the *dr_identifier* is encoded as a string and directed to the *General-Purpose Compressor*.

#### Semi-Duplication Removal

The *Semi-Duplication Removal* block finds similar, but not identical, nucleotide sequences, and groups the sequences, before encoding and removing the similar part of the sequences within each group in an efficient manner to reduce the compression ratio (example of workflow in **Fig. 4**). In this context, two sequences are considered semi-identical if their initial non-identical character fall within a certain number of characters from their respective endings, defined as the *sdr_threshold*. While the *Semi-Duplication Removal* block focuses on similarities at the beginning of the sequence, there can also be similarities at the end of the sequence. To ensure removal of similar sequences at the end of a sequence, the GeneSqueeze algorithm sends sequences to the Sequence Reversion block prior to inputting the sequences into another instance of the *SDR* block.

**Figure 4.**
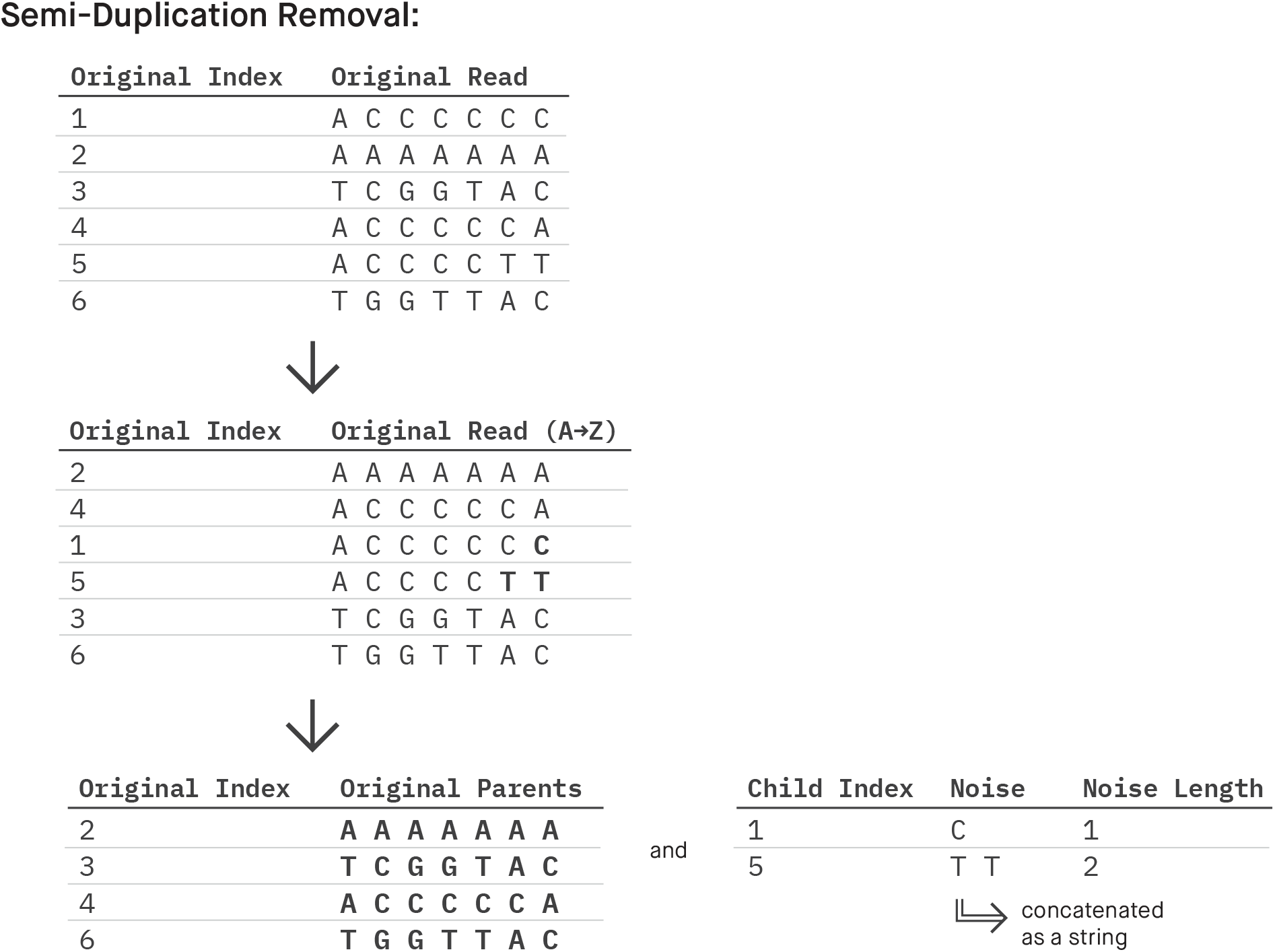
GeneSqueeze Semi-duplication removal process

In *SDR* operation, the sequences are first ordered alphabetically with their original indices being tracked. Once sorted, semi-duplicate nucleotide sequences are referred to as groups. For each group of nucleotide sequences, only the first instance of the sequence is designated as the parent sequence. The remaining semi-duplicate sequences within the group are designated as the *child sequences*. Each child sequence is assigned a *child index* that indicates which parent each child sequence belongs to. GeneSqueeze examines the non-identical nucleotides between two sequences. We specifically encode the subsequence beginning from the first non-identical character and extending to the last character of the sequence which we define as “noise”. The algorithm identifies noise between the first child and its parent and between any subsequent child with its preceding child. The length of the “noise sequence” is also encoded as “*noise length*”. Noise length along with child identifier indices are passed to the *General-Purpose Compressor* (*noise length, child index*). Meanwhile, the *noise sequences* are directed to *Nucleotide Sequence Regulator* block. Finally, the *parent sequences* are resorted using their original indices and directed to *Sequence Reversion* or *Nucleotide Sequence Regulator*, if it is the first or the second *SDR* operation, respectively.

The pseudocode presented below shows the semi-duplication removal process:

FUNCTION semi_duplication_removal(sequences, sdr_threshold)

1. Convert sequences to an array
2. Sort the sequences and keep track of their original indices
3. Calculate the differences between adjacent sorted sequences
4. Prepare data structure to store information about sequences, groups, noise sequence, and noise lengths
5. Create groups of sequences based on the cut points
6. Identify the first sequence in each group as parent sequence and rest of the sequences as child sequences
7. Calculate the noises between each sequence and its previous sequence for each group
8. Calculate the length of obtained noises
9. Build a DataFrame to organize the information for the child indices
10. Sort sequences by original indices
11. Extract the indices of the child sequences as child indices
12. Join noise sequences as a single string
13. Return the child indices, noise lengths, noise sequences, and the parent sequences

#### Sequence Reversion

As mentioned in the *Semi-Duplication Removal* block, this algorithm is crafted to identify and eliminate sequences that exhibit semi-duplication either at their beginning or end. As the SDR block only identifies duplicates at the beginning of a sequence, we have to reverse the sequence to remove duplicates found at the end of the sequence. These reversed sequences are then sent through the *SDR* block, depicted in **Fig. 2**. After passing through SDR, the output parent sequences are reverted to their original orientation using the *Sequence Reversion* block.

#### k-mer Dictionary Optimizer

*k*-mers denote subsequences of length ‘*k*’ extracted from longer nucleotide sequences. *k*-mers serve as ideal units for partitioning nucleotides to enhance algorithmic efficiency. The frequently occurring *k*-mers can be identified, assigned short indices, and subsequently eliminated to conserve space. Each selected *k*-mer and its associated index is stored as a key-value pair in a dictionary. There exists a trade-off between the length and frequency of *k*-mers. The elimination of longer *k*-mers can result in larger deletions overall, but only if these *k*-mers are sufficiently high in frequency. Removing shorter *k*-mers results in removing less information per sequence; however, given their higher likelihood of occurrence, removal of high frequency short *k*-mers can lead to a greater number of nucleotides being eliminated.

This dichotomy indicates that it is imperative to ascertain the optimal *k*-mers to maximize the compression ratio. As there are many k-mers within a dataset we can store the optimal *k*-mers in a *k*-mer dictionary which can then be used for compression. The *k*-mer dictionary can contain many *k*-mers of numerous *k* lengths; given the myriad of combinations, the creation and storage of this dictionary can be an expensive process. To enhance cost effectiveness of the algorithm, GeneSqueeze creates an initial *k*-mer dictionary using a small random subset of nucleotide sequences, assuming similar distribution of *k*-mers across the whole dataset. The created *k*-mer dictionary can then be used to reduce redundancy across all remaining nucleotide sequences.

In theory, the storage space required for both the *k*-mer dictionary and the additional encoding information after *k*-mer removal can exceed the space saved by removing *k*-mers. A potentially confounding factor is that the removal of each *k*-mer from the remaining sequences may affect the frequency of other *k*-mers being found within the sequences. Thus, we need an algorithm to iteratively find the ‘best’ remaining *k*-mer and assess the worth of adding that k-mer to the dictionary, considering both space savings and encoding costs. Therefore, a dictionary with both an optimal count of *k*-mer – index pairs and of *k*-mers of optimal sizes is desirable and thus these two optimization problems are inherently coupled together and require simultaneous computation. The optimal *k*-mer dictionary is specific to each sample’s sequences, and can vary significantly between samples. Thus, in this block, an optimization strategy is presented to solve the optimization problems and find an optimal *k*-mer dictionary for each sample so that in *k-mer Removal Block we* encode the *k*-mers for each sample using its ad-hoc *k*-mer dictionary (**Fig. 5**).

**Figure 5.**
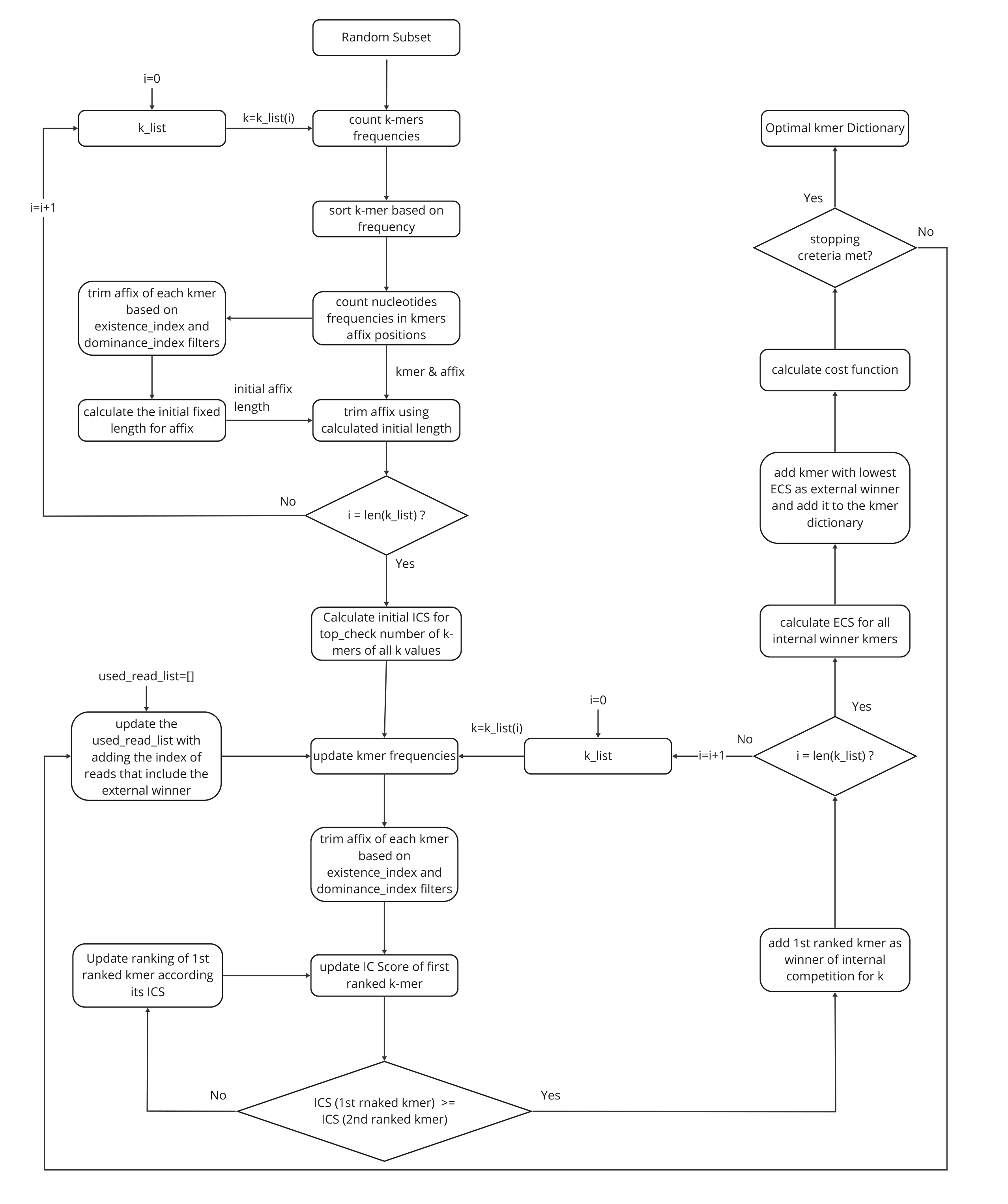
GeneSqueeze *k*-mer dictionary optimizer process

Given the fact that nucleotide sequences will align to specific locations in a reference genome, they are not inherently randomly distributed. This indicates there is a high level of imbalance for the substrings prior to or next to each *k*-mer, which is exploited to achieve higher levels of compression in our algorithm. GeneSqueeze finds high frequency *k*-mers and filters their prefix and/or suffix based on given thresholds. During encoding, it looks for an exact match for the selected optimal *k*-mers and then checks if the substrings before and after them match with the identified prefix and suffix, respectively. If not, the difference between them is encoded as noise.

Among the hyperparameters of this algorithm, the *k* values for each *k*-mer in the *k*-mer dictionary play a significant role. These *k* values are not limited to a single number but can consist of multiple values. A pre-selected list of *k* values, referred to as *k_list*, is considered for the *k*-mer selection process. As previously noted, there exists a trade-off between the length and frequency of *k*-mers, making it crucial to ascertain the optimal *k*-mers. In deciding which *k*-mer is more qualified to be first added to the *k*-mer dictionary, the algorithm takes into account the storage saved by removing the *k*-mer from sequences. A simplistic example which includes a small *k_list* and is without affix: if *k_list=[k1, k2]* where *k2 > k1*, and if frequency of a selected *k2*-mer is higher than a selected *k1*-mer, the selection of k2 will lead to larger improvements in compression ratio. Conversely, if the frequency of the selected *k2*-mer is lower than that of the selected *k1*-mer, a trade-off between their length and frequency is taken into account. The algorithm outlined in this section streamlines the process of selecting *k*-mers using these calculations.

As can be seen in the diagram of **Fig. 5**, for every *k* value within *k_list*, internal competition arises among the *k*-mers at each iteration. Subsequently, winners of these internal competitions, one for each *k* in *k_list*, engage in an external competition to determine the overall external winner. The victor of the external competition joins the optimal *k*-mer dictionary, while the external losers revert to their respective internal competitions. The internal and external competitions are done based on internal competition score (*ICS*) and external competition score (*ECS*), and are presented in equations 4 and 5, respectively.

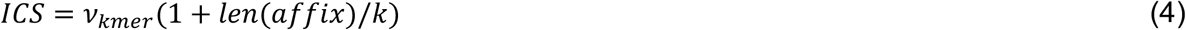

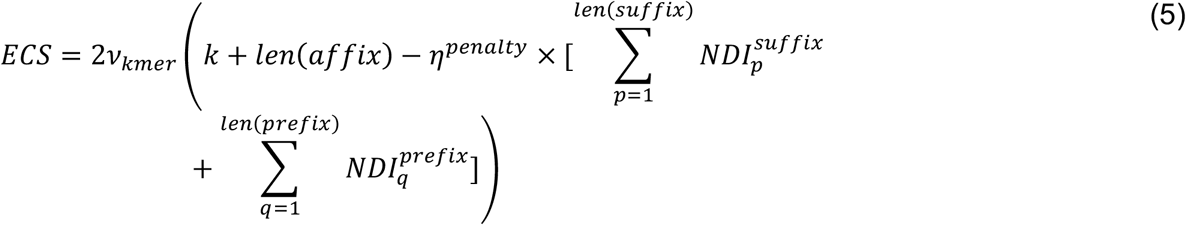

In this study, *ν* represent the frequency of *k*-mer. Furthermore, *η*^*penalty*^ is the penalty factor for the fraction of nucleotides that does not match the prefix-suffix. This penalty factor helps to include the cost of prefix-suffix mismatch in calculation of *ECS* and the cost function presented in equation 9. It is imperative during encoding, as elucidated in the *k*-mer removal block, to store not only the correct type of nucleotides but also the position and quantity of noises. As is shown in equation 6, the existence index for position *i* (*i*^*th*^ character) of affix, *EI*_*i*_, is defined as the total frequency of all bases in position *i, ω*_*i*_, over the frequency of the *k*-mer itself, *ν*.

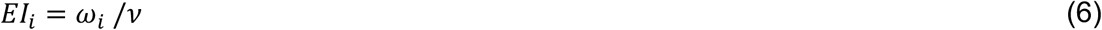

Also, as indicated in equation 7, the dominance index *DI* for position *i* of affix is the frequency of the dominant base position *i, λ*_*i*_, over the total frequency of all bases in position *i* of the *k*-mer affix, *ω*_*i*_.

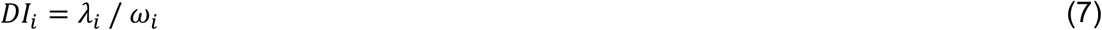

The non-dominance index *NDI* for position *i* of affix is presented in equation 8.

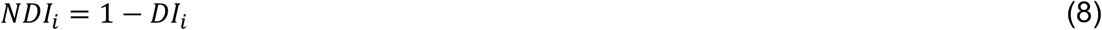

An example for the prefix-suffix identification process, including calculation of the dominance and existence indices, is presented in **Fig. 6**. *EI_threshold* and *DI_threshold* are the considered thresholds for *EI* and *DI*, respectively. As depicted in **Fig. 6**., the frequency of the dominant nucleotide at each specific position before and after the selected *k*-mer is shown in blue. Among these, the green nucleotides surpass *EI* and *DI* thresholds (*EI_threshold* and *DI_threshold*) and are selected as the *k*-mer prefix and suffix.

**Figure 6.**
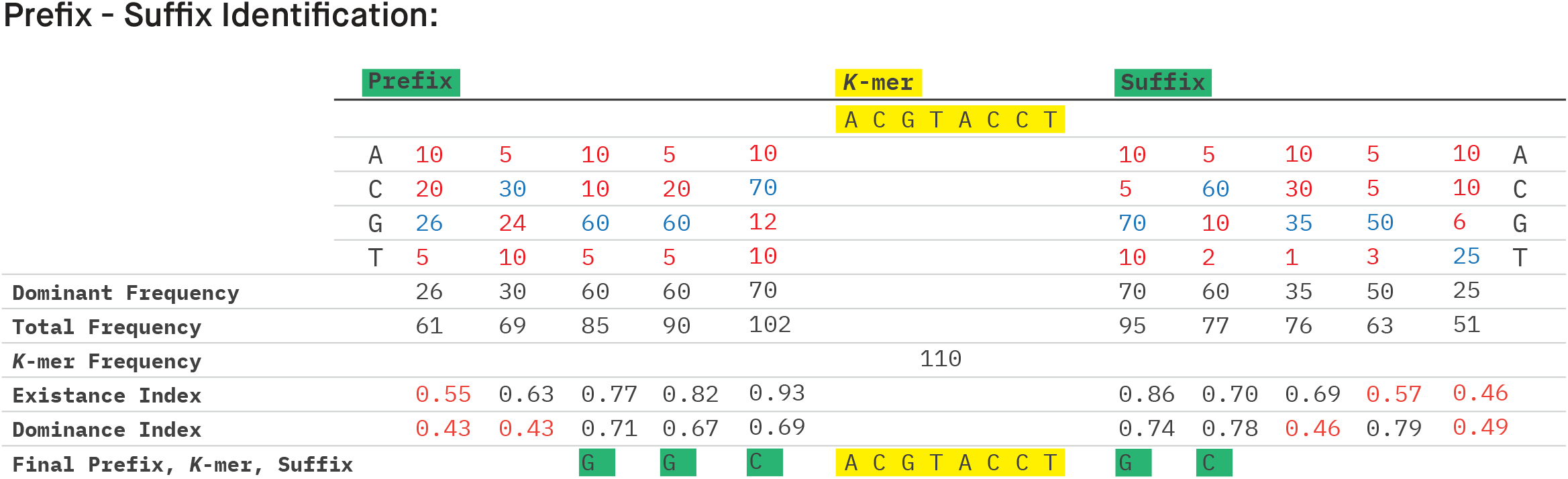
GeneSqueeze Prefix-suffix identification process for *EI_threshold = 0*.*6* and *DI_threshold = 0*.*5*

This process persists until one of the following stopping criteria is fulfilled:

1. The number of consecutive cycles with an increase in cost function score exceeds the “tolerance_threshold.” The algorithm allows for temporary increases in the cost function score over a set number of cycles, denoted as “tolerance_threshold”. If the number of cycles proceeds beyond the set threshold then the process is stopped. The algorithm then removes the last *tolerance_threshold* elements (*i*.*e*. the last *k*-mer that was added to the *k*-mer dictionary) added to the dictionary, and passes the finalized dictionary to the subsequent stage. The *tolerance_threshold* value remains constant throughout the process.
2. The length of the *k*-mer dictionary reaches the *max_dict_len* limit.

In this study, the “data size” is denoted by *β*. By using the following formulas (equations 9-19), which have been derived based on the GeneSqueeze algorithm, without compressing the data, we can have a rough estimation of *β*_*compressed_seqs*_ in bits as a function of *dict_len* for the existing distribution of *k*-mers in the FASTQ/A file. *dict_len* denotes the number of *k*-mers added to the dictionary. To calculate *β*_*compressed_seqs*_ in equation 9, *β*_*original_Seqs*_ is calculated based on equation 10, *β*_*kmer_all*_ based on equations 11 and 13-17, and *β*_*affix_all*_ based on equations 12, 18 and 19. In equation 9, it is expected that *β*_*kmer_all*_ + *β*_*affix_all*_ will be negative. As such, it follows that *k*-mer removal is a better option than simply computing binary encoding on all nucleotide sequences. It is worth noting that in the cost function, which is designed to provide an approximate estimation of actual compressed data, all bases are assumed to be regular, forming the basis of equation 10. In this equation *seq_depth* and *seq_length* are the number and length of the input nucleotide sequences of this block, respectively.

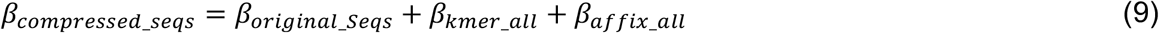

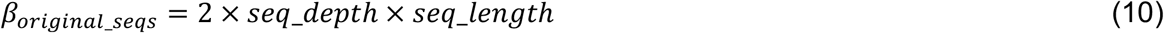

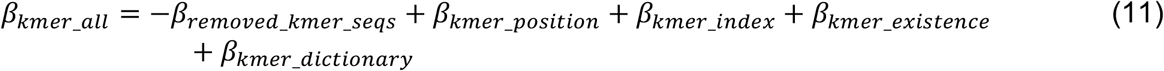

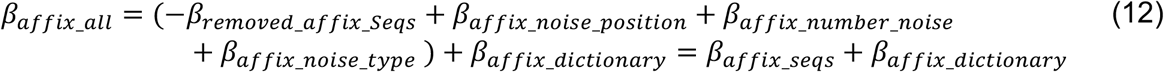

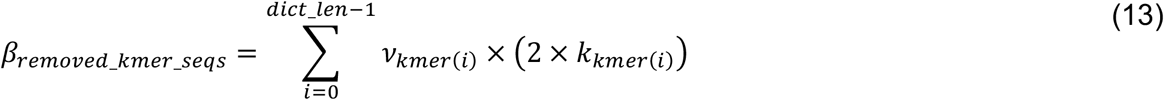

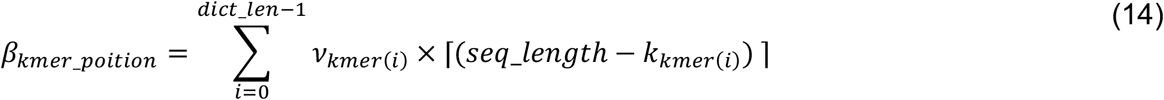

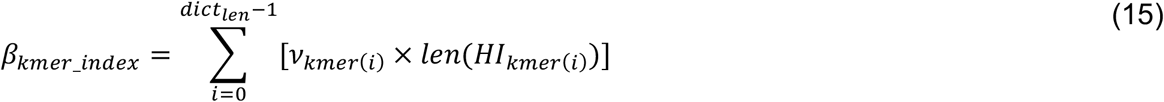

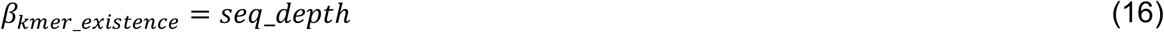

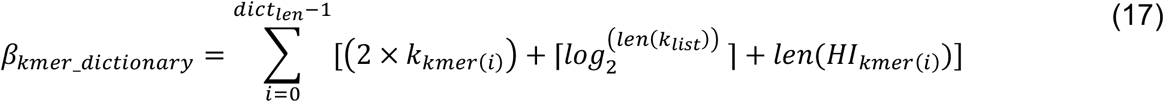

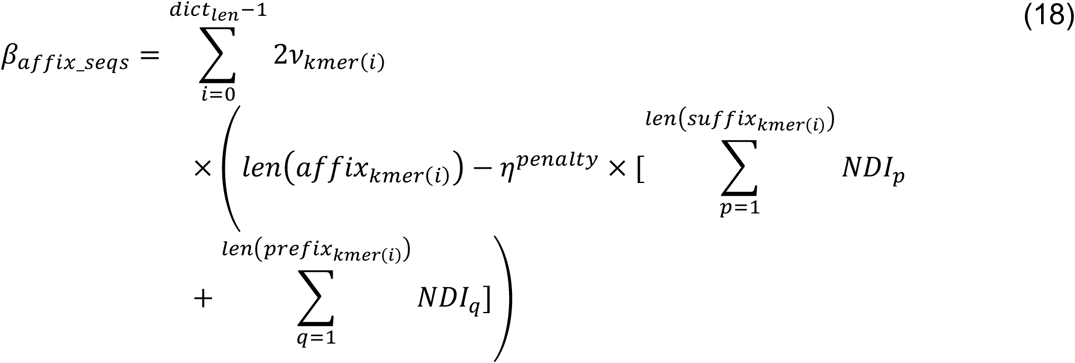

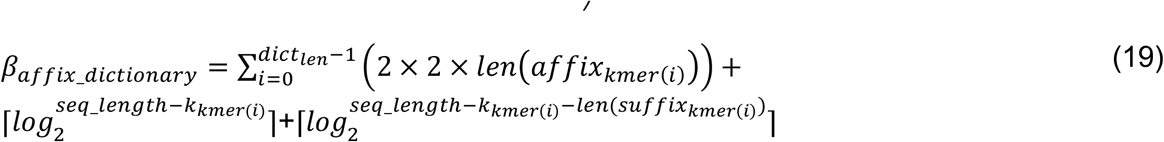

where, *HI* stands for binary code length of Huffman encoded *k*-mer in *k*-mer dictionary. *Existence* is a binary vector of length *seq_depth* to show which nucleotide sequence include any *k-*mer of the final dictionary.

The dictionary optimizer obtains the optimum *dict_len* value to have the *β*_*compressed_seqs*_ minimized.

#### Random Sequence Selector

As we explained, *k*-mer dictionary creation is an expensive process. To reduce computational load we utilized the *Random Sequence Selector block*, which randomly selects a subset of nucleotide sequences to be used in *k-mer Dictionary Optimizer* block. The percentage of sequences selected is defined by *random_subset %*.

#### k-mer Removal

The *k*-mer removal block removes duplicative *k*-mers in the dataset. The function first checks each nucleotide sequence of the input sequences to determine if there is a full match for the *k*-mers in the finalized dictionary. If there is a match, it removes the *k*-mer and its affix and encodes its existence (one bit per nucleotide sequence), *k*-mer index, *k*-mer position in nucleotide sequence and its index (dictionary value), as well as the affix noises. After removing the *k*-mer and its affix, the subsequences before and after the removed part of the sequence are concatenated and sent to the *Nucleotide Sequence Regulator* block. The untouched nucleotide sequences (nucleotide sequences with no identified *k*-mers) are passed to the *Semi-Duplication Removal* block. The process is shown in **Fig. 7**.

**Figure 7.**
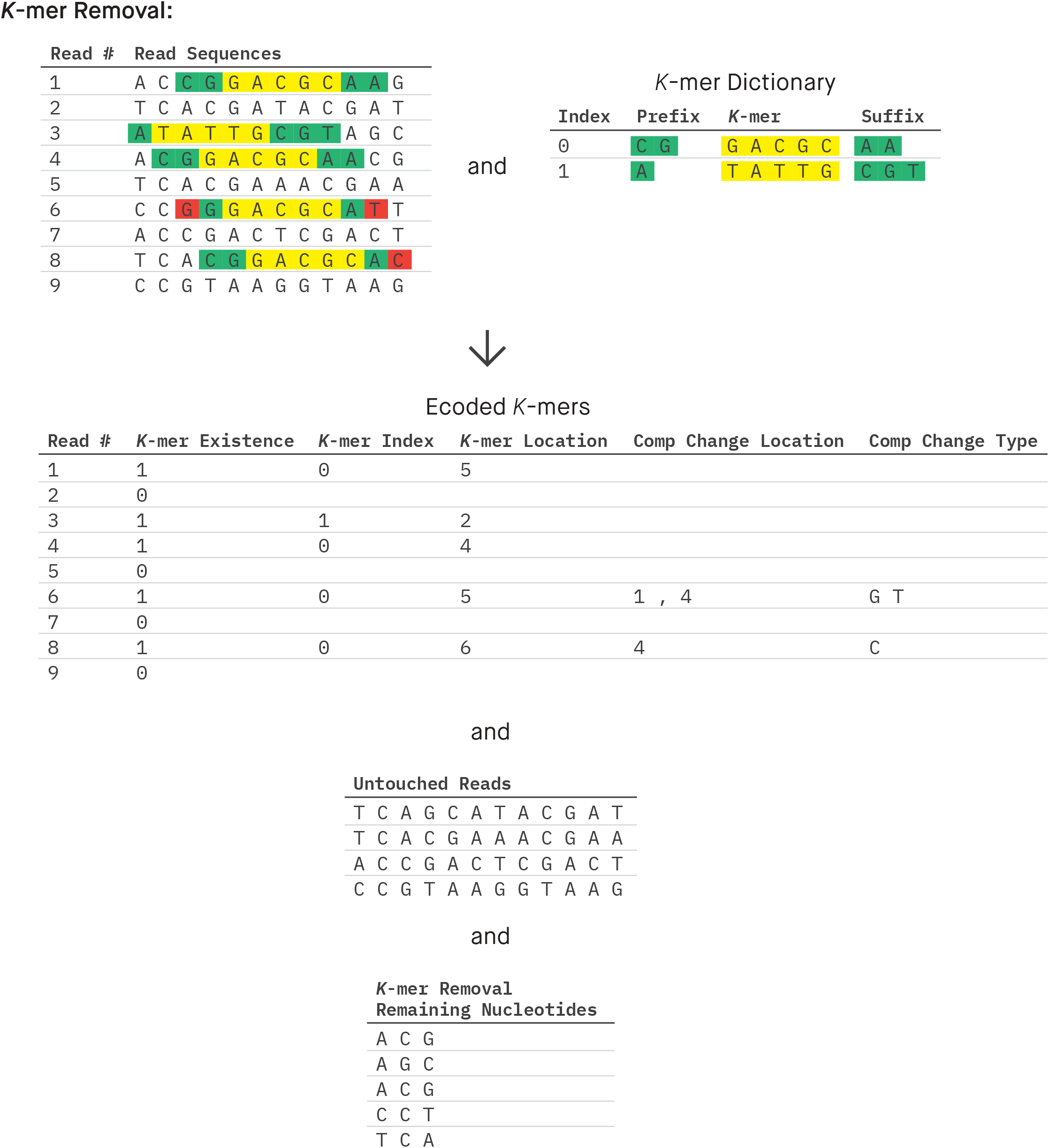
GeneSqueeze *k*-mer removal process

#### Nucleotide Sequence Regulator

This block encodes any bases presented in **Table 1** other than A, C, G and T. As is shown in **Fig. 2**, the input for this block is the output of all other blocks that are in the nucleotide sequence form. Given that the predominant characters in nucleotide sequences are usually A, C, G, and T, and considering that four characters can be encoded using only two bits in binary, this block encodes the information of non-ACGT characters separately and then replaces them with A in the nucleotide sequence. Therefore, the output of this block is the nucleotide sequences that only include A, C, G and T, as well as the position of non-ACGT characters. They are directed to *Nucleotide Sequence Binary Encoding* and *General-Purpose Compressor* blocks, respectively.

This block assumes by default that all irregular (non-ACGT) nucleotides characters are N (which is the most common irregular nucleotide) and encodes their nucleotide sequence position. In cases where any irregular non-N nucleotide is observed, as well as the position, its actual character is also encoded.

#### Nucleotide Sequence Binary Encoding

The nucleotide sequences are then encoded in binary using the hash table presented in **Table 2**. As mentioned in the *Nucleotide Sequence Regulator* block, irregular nucleotides are encoded as A: [0 0] since their information is recorded separately.

**Table 2.** Nucleotides binary encoding hash table.

| Nucleotide | Binary code |
| --- | --- |
| A | 00 |
| C | 01 |
| G | 10 |
| T | 11 |

#### Quality Score Compression Branch

The segmented quality scores are passed to the quality score compression branch of the algorithm, which has one block dedicated to duplication removal.

#### Duplication Removal (Quality Score)

After quality scores are segmented in *Sequence Segmenter* block, they proceed through the *Duplication Removal* block. This block searches across the quality score sequences to find and group identical sequences and encodes only the first instance of a specific quality score sequence of each group, designated as the parent sequence. The other sequences of each group are designated as child sequences and are referred back to the parent in the same fashion as nucleotide sequences. The encoded files are sent to the *General-Purpose Compressor*.

#### Read Identifier Compression Branch

The segmented read identifiers are passed through the read identifier compression branch of the algorithm which has one block in which the read identifiers are encoded.

#### Read Identifier Encoder

The *Read Identifier Encoder* block stores the first read identifier. The block then identifies which portions of the identifier are fixed or variable and subsequently encodes only the change in variables for each ensuing read.

#### General-Purpose Compressor

All the generated files are then sent to a general-purpose compressor to achieve a higher compression ratio. GeneSqueeze uses BSC, which is a general-purpose compressor built based on the Burrows-Wheeler Transform (BWT).^24^

## Results and Discussion

We conducted a case study where we compared GeneSqueeze against two other algorithms for the compression of nucleotide-containing FASTQ files: (1) the prevalent general-purpose compressor, gzip (version 1.9) and (2) the domain-specific compressor, SPRING (version 1.1.1). ^25^

### Dataset

We tested the performance of each compression algorithm on a collection of datasets that totaled 1572 GB in uncompressed size. It contained 283 FASTQ files, ranging in length (**Table 3**), depth (**Table 4**), and size (**Table 5**). The full list of the datasets used for experiments is found in **Table S1**. In these experiments, each FASTQ file is compressed independently. The characteristics of each dataset can directly affect the performance of a compression algorithm, thus algorithm performance was analyzed by characteristics. Natively, these datasets contained N but no non-N irregular nucleotides. So, to test the losslessness of non-N irregular IUPAC nucleotides, two N characters were replaced with non-ACGTN IUPAC characters in a FASTQ file. Additionally, to alleviate memory consumption, four 50+ GB files were split into four chunks each during GeneSqueeze encoding and recombined after decoding.

**Table 3.** FASTQ file size (GB) of the datasets used for experiments.

| FASTQ File Size (GB) | Number of Files |
| --- | --- |
| 0-4 | 156 |
| 4-8 | 75 |
| 8-12 | 48 |
| 54-57 | 4 |

**Table 4.** Read depth (million reads per sample) of the datasets used for experiments.

| Read Depth (million reads per sample) | Number of Files |
| --- | --- |
| 1-20 | 172 |
| 20-35 | 40 |
| 35-65 | 67 |
| 215-224 | 4 |

**Table 5.** Read length (nucleotide sequence) of the datasets used for experiments.

| Read Length (nucleotide sequence) | Number of Files |
| --- | --- |
| 35-36 | 106 |
| 50-51 | 107 |
| 100-101 | 63 |
| 202 | 7 |

### Hyperparameters

The values of the hyperparameters for GeneSqueeze are presented in **Table 6**. GeneSqueeze’s *k-mer Dictionary Optimizer* block selected the optimal length of the *k*-mer dictionary and the *k*-mers via the optimization process presented in the Methods section (*k-mer Dictionary Optimization* block). In the current version of GeneSqueeze, we employed a static value for the rest of the hyperparameters, which is not optimal. Our future directions include upgrading the algorithm to dynamically determine and adapt the optimal values of the hyperparameters for each dataset or sample to further improve our compression ratio.

**Table 6.** Hyperparameter values used by the GeneSqueeze algorithm.

| Hyperparameter | Value |
| --- | --- |
| k_list | [10, 25] |
| random_subset | 1% |
| EI_threshold | 0.1 |
| DI_threshold | 0.5 |
| tolerance_threshold | 10 |
| max_dict_len | 64 |
| top_check | 10 |
| $\eta^{penalty}$ | 6 |
| *ideal_len* | 80 |
| sdr_threshold (1st SDR block) | 5 |
| sdr_threshold (2nd SDR block) | 10 |

### Evaluation metrics

We measured the performance of GeneSqueeze against gzip and SPRING using the evaluation metrics presented in equations 20 and 21:

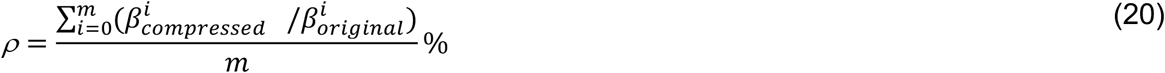

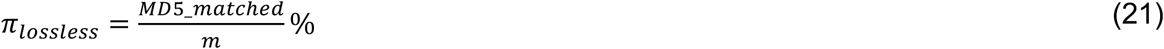

where *ρ* and *Π*_*lossless*_ are *average compression ratio (%)* and *losslessness ratio (%)*, respectively. Here, *m* represents the total number of FASTQ files in the test set, while *MD5_matched* denotes the number of samples with matched MD5 sums. The metric compression ratio was chosen to assess the total reduction in file size after compression, and losslessness ratio was chosen to understand if total data integrity was preserved after compression. These represent important factors in the overall efficiency and integrity of each compression algorithm.

### Compression results

All three algorithms were able to significantly compress the datasets. The average compression ratio of SPRING and GeneSqueeze were similar (7.61% vs 7.70%), and both were significantly lower than gzip (21.20%) (**Table 7**). FASTQ files can vary in size, read depth and read length, characteristics which we hypothesized could lead to changes in algorithmic efficiency. To understand which FASTQ files GeneSqueeze excelled at compressing, we examined compression ratio amongst FASTQ files with differing dataset sizes (**Table 8**), read depths (**Table 9**), and read lengths (**Table 10**). We found that both GeneSqueeze and SPRING outperformed gzip in all categories. We found that GeneSqueeze compressed files of size 8-12GB most efficiently and was least efficient at compressing large files (54-57 GB). Notably, GeneSqueeze compressed smaller files (0-4GB) better than SPRING (**Table 8**). When examining GeneSqueeze performance for differing read depths we found that GeneSqueeze was best at compressing files with read depths of 35-65 million reads and performed worst when compressing files with large read depths of 215-224 million reads. GeneSqueeze outperformed SPRING on read depths of 1-20 and 20-35 million reads (**Table 9**). Next, we examined compression ratio based on read length. We found that GeneSqueeze had the greatest compression on files with read lengths of 50-51 nucleotides and least compression on reads of 100-101 nucleotides. GeneSqueeze outperformed SPRING on files with read lengths of 35-36, 50-51, and 202 nucleotides (**Table 10**).

**Table 7.** Average compression ratio (%) of samples uncompressed and compressed with gzip, SPRING, or GeneSqueeze compression algorithms.

| Uncompressed | Compressed |  |  |
| --- | --- | --- | --- |
|  | gzip | SPRING | GeneSqueeze |
| 100% | 21.20% | 7.61% | 7.70% |

**Table 8.**
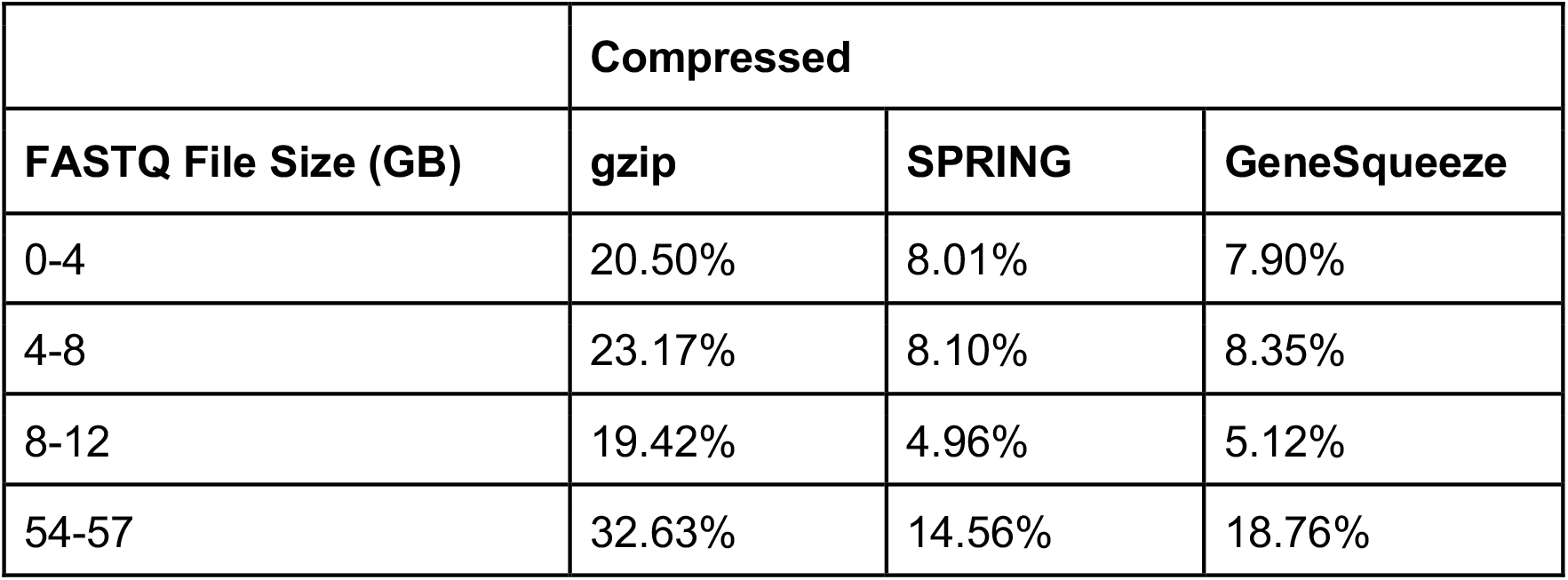
Average compression ratio (%) of samples by FASTQ file size (GB).

**Table 9.**
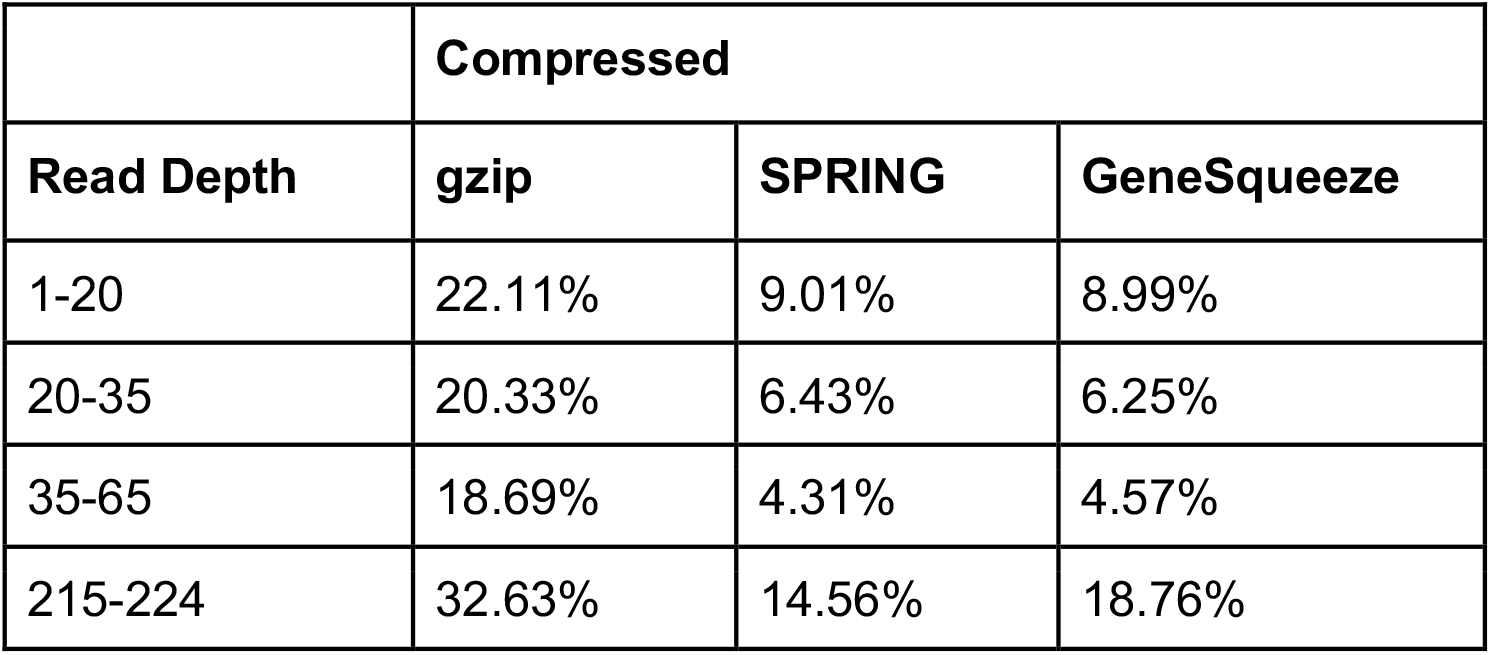
Average compression ratio (%) of samples by read depth (million reads per sample).

|  | **Compressed** |  |  |
| --- | --- | --- | --- |
| **Read Depth** | **gzip** | **SPRING** | **GeneSqueeze** |
| 1-20 | 22.11% | 9.01% | 8.99% |
| 20-35 | 20.33% | 6.43% | 6.25% |
| 35-65 | 18.69% | 4.31% | 4.57% |
| 215-224 | 32.63% | 14.56% | 18.76% |

**Table 10.** Average compression ratio (%) of samples by read length.

|  | **Compressed** |  |  |
| --- | --- | --- | --- |
| **Read Length** | **gzip** | **SPRING** | **GeneSqueeze** |
| 35-36 | 20.59% | 7.69% | 7.56% |
| 50-51 | 17.82% | 5.36% | 5.25% |
| 100-101 | 27.16% | 10.60% | 11.64% |
| 202 | 28.39% | 13.93% | 11.62% |

Quality scores are typically more challenging to compress than nucleotide sequences.^3^ Thus, quality score compression represents an area to be improved in the field. GeneSqueeze uniquely pre-compresses quality scores with a domain-specific compression process and then feeds that to the general-purpose compressor. Our method allows us to further reduce the file size dedicated to quality scores without any loss or impact on downstream analysis. The pie charts in **Fig. 8** indicate how much - on average - each component of a FASTQ file contributes to the total size, before and after GeneSqueeze compression. The difficulty in compressing quality scores due to its aforementioned characteristics is indicated by 60% of GeneSqueezed FASTQ file sizes, on average, being allotted to quality scores (**Fig. 8**). We observed that read identifiers were the easiest to compress due to their fixed template.

**Figure 8.**
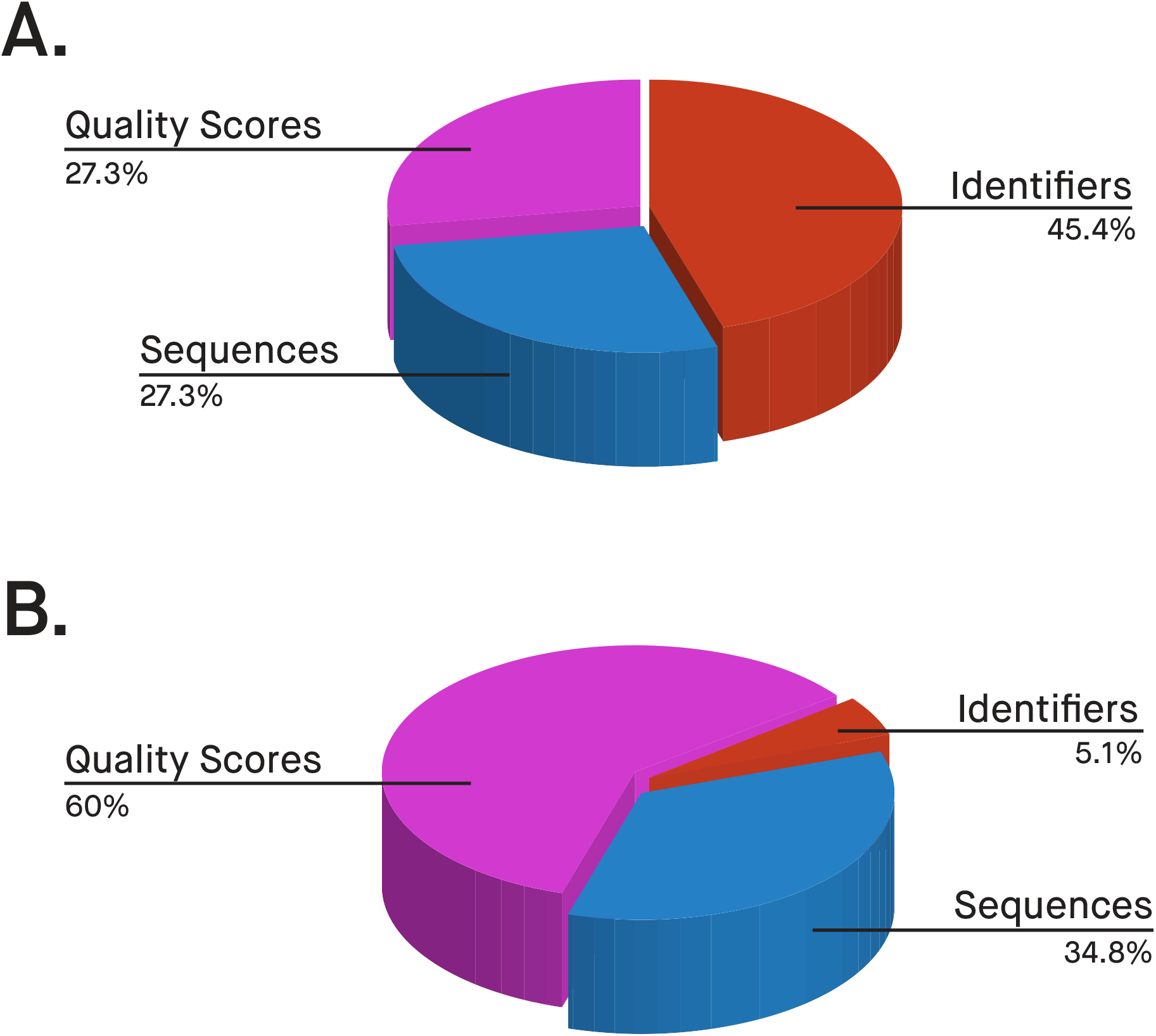
Average contribution of each of three parts of FASTQ files in a) uncompressed and b) GeneSqueezed files

We have also performed algorithm development and implementation for decompression of GeneSqueezed FASTQ files. To examine data accuracy and integrity of each compression method, we generated MD5 checksums for the original and decoded FASTQ files for all 283 files in the datasets. We observed 100% losslessness for GeneSqueeze and gzip, with 283/283 files having an MD5 match. However, SPRING experienced some loss during decoding.

SPRING had an MD5 match for 10/283 files for 3.5% losslessness. As has been mentioned in the SPRING Supplementary Data, SPRING discards any metadata or comments that follow the “+” in the second read identifier line (the line after nucleotide sequence and before quality score). ^14^

As our datasets did not include non-N irregularities, we changed two nucleotides in a FASTQ file to non-N irregularities in a separate experiment. We observed that GeneSqueeze and gzip losslessly encoded and decoded the file and that the MD5 hashes for the original and decoded files matched. However, SPRING was not able to decode non-ACGTN letters that exist in the IUPAC nucleotide code, meaning it was not IUPAC lossless. Overall, GeneSqueeze outperformed competitors by compressing file sizes with a compression ratio on par with that of current state-of-the-art compressors, all while maintaining losslessness.

## Conclusion

Nucleotide sequencing data contains intricate patterns of redundancy and variation that require specialized data compression techniques. The unique features of nucleotide sequences, such as hierarchical structure and redundancy, can be exploited to achieve high compression ratios while minimizing the loss of information. Developing effective compression algorithms for sequencing data requires a deep understanding of the data and the biological processes that generate it. The GeneSqueeze algorithm is capable of compressing FASTQ and FASTA data containing nucleotide sequences and is designed to be lossless for all parts of the FASTQ format, including the read identifier, quality score, and nucleotide sequence. GeneSqueeze relies on reducing the dimension and redundancy in genomic data in a unique and optimal way, and prior to storage in binary format.

The results of our comparison demonstrate the effectiveness and efficiency of our novel data compressor, with high and accurate data compression in our experiments. GeneSqueeze implements a number of functionalities that not only enhance compression ratios, but allow it to handle sequencing data losslessly.

This algorithm’s unique key features are:

1. Presenting a dynamic ad-hoc protocol for optimally encoding high frequency *k*-mers
2. Exploiting the redundancy in quality scores before passing them to the general compressor
3. Encoding and losslessly decoding all IUPAC characters
4. Expressing no limitation in read length / depth, allowing for flexibility in long-read / high-throughput FASTQ/As compression.
5. Compatible with all FASTQ/As identifier formats.

Overall, we observed that GeneSqueeze performed competitively against SPRING, with both performing better than gzip for compression ratios. GeneSqueeze and gzip maintained losslessness, whereas SPRING experienced loss in the majority of samples, based on MD5 matches. Due to GeneSqueeze’s novel approach, competitive performance, and complete preservation of genomic data, we believe GeneSqueeze is able to support the growing application of omics technologies in biomedical or clinical research. Our forthcoming emphasis will be on implementing GeneSqueeze in a more efficient programming language and in a manner that optimizes memory usage. GeneSqueeze’s current Python implementation reveals certain drawbacks in speed and memory usage when juxtaposed with SPRING and gzip, which are predominantly implemented in C/C++. Additionally, it is imperative to upgrade the algorithm to dynamically optimize the values of hyperparameters across multiple blocks, ensuring optimal adaptation for each dataset or sample.

## Acknowledgments

The authors would like to express their gratitude to individuals from Rajant Health Incorporated for valuable discussions and technical contributions, and to Rajant Corporation for providing funding support for this project.

The SPRING software used by the authors, was developed at the University of Illinois at Urbana-Champaign and Stanford University.

## Author contributions

Algorithm development: F.N.

Implementation: F.N., S.P.

Figure Generation: F.N., M.L.

Pseudocode Generation: F.N., R.C.

Conceptualization: F.N., S.P., E.K.M., G.S.

Manuscript Writing: F.N., M.L., R.C., E.K.M., G.S.

Review: F.N., S.P., E.K.M., G.S., M.L. R.C.

All authors revised and approved the manuscript.

## Data Availability

Datasets are sourced from the public databases listed in the Supplementary data.

